# Acoustic chip for rapid label-free early-stage detection of rare leukemic cells

**DOI:** 10.1101/2024.06.03.597122

**Authors:** Bruno F.E. Matarèse, Andrew Flewitt, Brian J.P. Huntly

## Abstract

We present a promising approach for detecting as few as 1% of rare leukemia cells during the early stages of blood cancers. This study demonstrates a novel microfluidic chip utilizing a bulk piezoelectric ceramic device to manipulate cells with sound waves. By analyzing the movement patterns of normal mononuclear cells (MNCs) and abnormal THP-1 acute myeloid leukemia (AML) cells within an acoustic field, we observed distinct behaviors. Our findings suggest a label-free, non-targeted approach for sensitive detection of rare abnormal cells within a mixed population. This method, based on acoustophoresis principles, holds promise for analyzing biophysical properties of individual cells for early cancer diagnosis, potentially leading to earlier intervention and improved patient outcomes for leukemia. While this study focuses on microscopic analysis, we also discuss the potential for developing large-scale acoustophoresis-based methods for high-throughput rare cell detection using high-resolution nanofabrication techniques.

## I. Introduction

Leukemia, a cancer of the blood-forming tissues, impairs the body’s ability to produce healthy blood cells. This disruption leads to an overproduction of abnormal white blood cells, hindering the immune system’s capacity to fight infection and the bone marrows production of red blood cells and platelets, the building blocks for blood clots. Early detection of leukemia is important for successful treatment and improved patient outcomes. Early detection of Acute Lymphoblastic Leukemia (ALL), a common type in children, can significantly increase the 5-year survival rate to over 90%. However, current diagnostic methods, such as complete blood counts (CBCs) and bone marrow biopsies, have limitations that render early detection difficult [1]. While bone marrow biopsies offer definitive diagnoses, they are invasive and can cause discomfort [2]. Complete blood counts (CBCs), on the other hand, often struggle to identify pre-malignant cells due to their rarity and the subtle changes they exhibit, such as altered morphology, variations in cell surface markers, or underlying chromosomal abnormalities that may not be readily detectable. Indeed, early-stage abnormal cells can be challenging to detect through blood tests due to the low number circulating at this stage [3]. Furthermore, obtaining biopsy results can take several days, potentially delaying treatment initiation [4]. These limitations highlight the critical need for innovative, non-invasive, and rapid tools for detecting early-stage leukemia or for those patients that require more sophisticated investigations [1].

This study explores the potential of using piezoelectric materials to generate acoustic standing waves with varying pressure regions (nodes and anti-nodes) within a microfluidic chip. Piezoelectric materials can convert electrical signals into sound waves and vice versa. Within the chip’s detection chamber (see Fig. 1.), these pressure variations may differentiate the movement of various cell types. This non-targeted approach holds promise for identifying potential abnormalities in cell movement, such as those indicative of early-stage leukemia. Different cell types often exhibit variations in their physical properties, for instance, in their stiffness [5]. By analyzing how cells move within an acoustic field, this method aims to detect subtle changes in these properties, potentially revealing signs of cellular dysfunction.

**Fig. 1.**
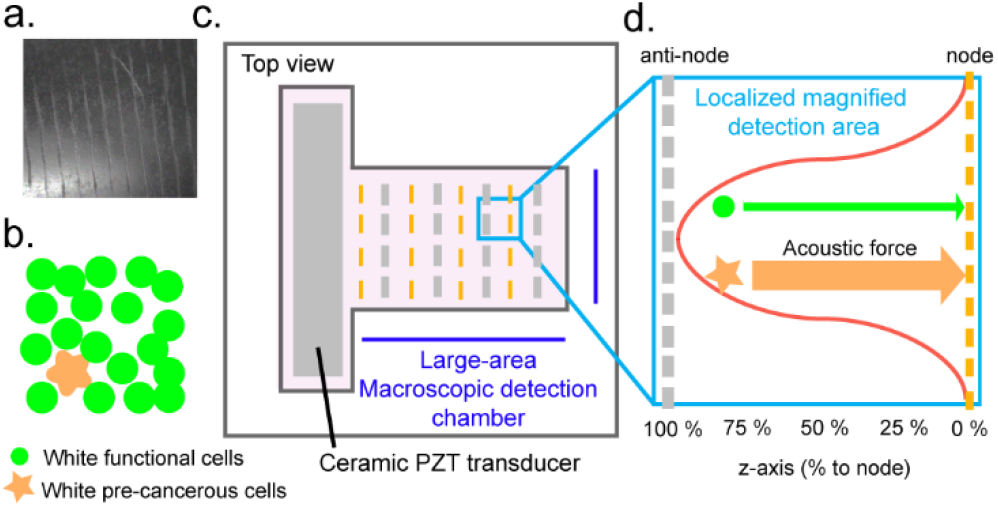
Label-free Differentiation of Leukemic Cells using Acoustic Chip. (a) Microscope image shows healthy white blood cells, formally called normal mononuclear cells (MNCs) aligned at specific points (pressure nodes) within the detection chamber. (b) Schematic depicts a mix of rare pre-cancerous cells (orange star) and healthy white blood cells (green circle). (c) Acoustic chip design with PZT transducer that creates sound waves and a large-area detection chamber for seeing the motion of many cells. The dashed gray lines show areas of high pressure (anti-nodes), and the dashed orange lines show areas of low pressure (pressure nodes). (d) Zoom-in localized magnification shows pre-cancerous cells (orange star) experiencing stronger forces (orange arrow) towards pressure nodes (dashed orange) meaning they move faster than healthy cells (green circle) that are subject to weaker forces (green arrow).

This novel, non-invasive method enables direct analysis of cells from blood samples, eliminating the need for invasive bone marrow biopsies. It offers a two advantages: analysis of intrinsic cellular properties (label-free) which preserves sample purity and compatibility with established biomarkers for downstream analysis. Furthermore, the method boasts a dramatic reduction in processing time, particularly for large samples. This approach leverages large-area macroscopic detection (as shown in Fig. 1.) to simultaneously observe individual cell motion for over 10,000 cells per acoustic enrichment run, which lasts less than 5 seconds.

This translates to analyzing cells thousands of times faster compared to the time-consuming process of cell-by-cell flow investigation. Importantly, the method remains efficient even for smaller detection areas analyzed with a single objective lens. In such cases, analyzing hundreds of cells still takes under an hour per million cells (over 95% faster than conventional methods). This rapid analysis is particularly advantageous for detecting subtle changes associated with early-stage leukemia, where well-defined biomarkers might be less prominent or indeed not known at all. In our initial study, we employ THP-1 cells, a well-established human acute myeloid leukemia cell line, due to their widely recognized characteristics and extensive use in leukemia research. Our initial findings suggest that this approach can easily differentiate between THP-1 leukemia cells and normal mononuclear cells (MNCs) (mononuclear cells, a heterogeneous collection of white cells present in the blood) based on their movement speed towards pressure minima (nodes) within the acoustic field. The success of this approach paves the way for a rapid, large-area detection chamber to facilitate the earlier identification of malignant blood cells, potentially enabling faster and more efficient patient care.

## II. Preparation and characterization

To investigate cell differentiation, peripheral blood mononuclear cells (MNCs) were isolated from healthy donors using standard density gradient centrifugation. In parallel, the THP-1 human AML cell line (ATCC® TIB-202™) was used as a model system for abnormal cells. To detect rare THP-1 leukemia cells, suspensions were prepared at 1 million cells/mL and mixed with healthy MNC suspensions (1 million cells/mL) at various ratios (0%, 1%, 5%, 10%, 100%). All samples were then diluted to a consistent density of about 20 cells per field of view (at 20x magnification) for analysis. This ensured fair comparison across samples, including controls with pure THP-1 and pure MNCs.

The microfluidic chip design incorporates two main components: a 10 mm × 10 mm microfluidic detection chamber and a separate 20 mm × 100 mm piezoelectric (PZT) chamber. These chambers are connected by an opening, allowing a well-defined standing wave to form within the resonant cavity of the detection chamber. Standard photolithography and soft lithography techniques were used to fabricate the chip with PDMS (polydimethylsiloxane). This process creates a rectangular microfluidic chamber with an open top, facilitating observation under a microscope, or with a closed top bonded to microscope slides. An open channel within the detection chamber allows the acoustic waves generated by the PZT element to efficiently interact with the cell-containing fluid. A 18 mm × 80 mm rectangular PZT (lead zirconate titanate) element of 2.1 mm thick is positioned within a separate chamber located beneath the microfluidic chamber. A Müller-Platte needle probe hydrophone measured the acoustic pressure distribution and intensity within the microfluidic channel. The C60 impedance analyzer (Cypher Instruments) identified an optimized resonance peak of the PZT at ~1 MHz. The operating frequency created a well-defined pressure pattern inside the chamber with alternating high (antinodes) and low (nodes) pressure points spaced half the wavelength (*λ*/2∼740 μm) apart. The pressure amplitude at the antinode reached 50 kPa, offering sufficient spatial variation for cell movement (without compromising viability [6]) and the achieved intensity (10-50 W/cm^2^) ensured effective cell manipulation towards the nodes.

To validate the method’s ability to detect rare cell populations, we initially focused on a smaller field of view within the microfluidic chip (see Fig. 1.d.). This approach was particularly suitable for detecting THP-1 leukemia cells, which comprise approximately 1% of the total cell population in our experiments. Five replicate measurements (*N*=5) were performed per sample, analyzing the same field of view within the detection chamber. To enable future implementation in large-area detection chambers, cell movement within the acoustic field (varying concentrations) was recorded using a high-speed camera (100 fps) mounted on an inverted Nikon Eclipse Ti2 microscope. Particle Image Velocimetry (PIV), a well-established optical method [7], was employed to analyze cell speed captured by the camera and processed by MATLAB and ImageJ. PIV decomposes the image into small interrogation windows and calculates the average displacement of particles within each window between consecutive frames. This displacement, divided by the time interval between frames (1/100 s at 100 fps), provides the average velocity of the particles within that window. To account for potential variations across the pressure gradient, the analysis focused on three distinct horizontal positions within the channel relative to pressure nodes and anti-nodes: 25%, 50%, and 75% of the node-to-antinode *z*-position length. These positions were identified beforehand using a multi-method approach (microbead manipulation, hydrophone characterization, and cell alignment visualization/post-processing).

## III. results and discussion

Ceramic piezoelectric transducers were used to investigate whether differentiation of THP-1 leukemia cells from normal MNCs could be achieved by utilizing an acoustic standing wave within a microfluidic chamber. This manipulation of cell movement revealed a trend where cells migrated differently towards pressure nodes, suggesting that variations in cell stiffness might influence their interaction with the acoustic field. Control groups containing only THP-1 cells or only MNCs were analyzed to understand how each cell type behaves under the sound waves in isolation. In order to understand how cells respond to the sound waves, we measured their speed at three specific locations within the pressure gradient (25%, 50%, and 75% of the distance between pressure nodes and antinodes) (see Fig. 2).

**Fig. 2.**
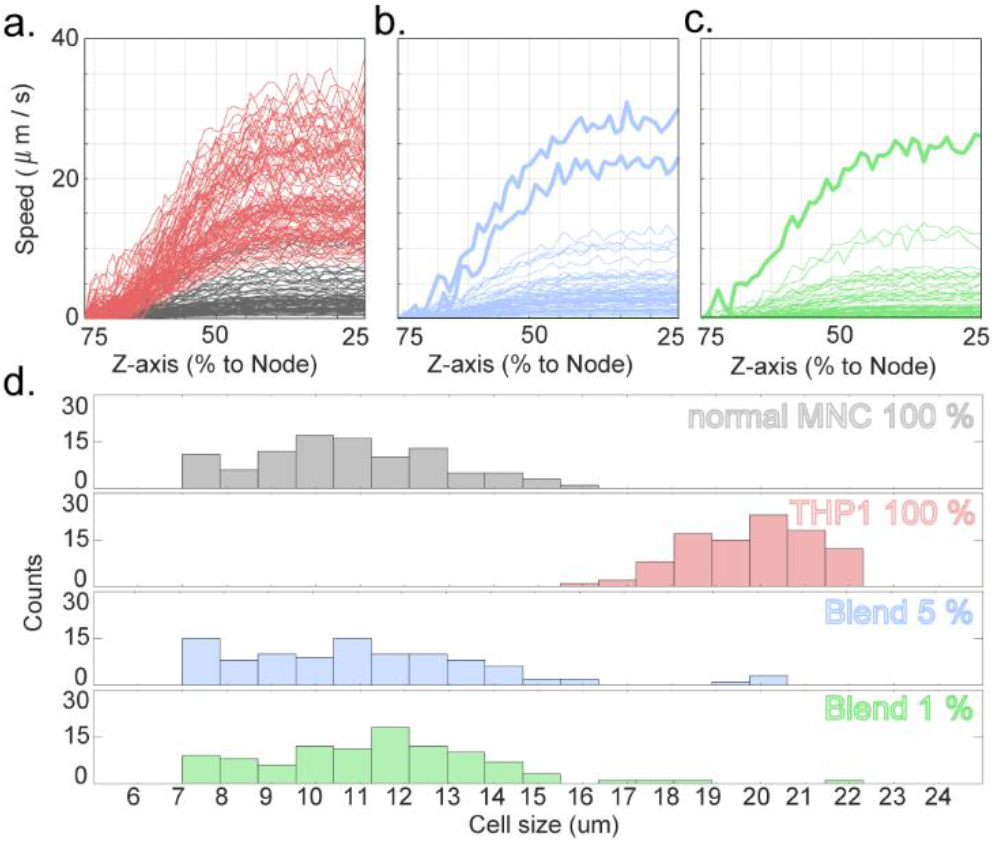
Cell Speed and Size Differentiation of THP-1 and MNCs under Pressure Gradient. Panels a-c) show average cell speed (μm/s) at various positions along the pressure gradient (***z***-axis) between 25% and 75% of the node-to-antinode distance (~100 cells investigated over *N*=5/condition). a) THP-1 cells (red) and normal MNCs (black); b) 5% THP-1 mixture with normal MNCs (blue) and c) 1% THP-1 mixture with normal MNCs (green). Panel d) shows the distribution of cell size (μm) for THP-1 cells (red), normal MNCs (black), 5% THP-1 mixture (blue), and 1% THP-1 mixture (green).

The analysis revealed that the pressure gradient affects cell speed. The clearest distinction between healthy MNCs and leukemic THP-1 cells occurred at about 25% from the pressure node (area with minimal pressure). This zone has less background noise, allowing us to better identify cell types based on how their speed changes across the pressure gradient. The analysis excluded areas very close to the nodes (<25%) and antinodes (>75%) for two reasons. First, these areas have minimal or maximal pressure changes, making measurements less accurate. Second, strong acoustic forces near these points cause cells to clump together, affecting their movement in the fluid. Analysis of both movement speeds and size distributions revealed a distinct characteristic that differentiates normal MNCs from THP-1 leukemia cells. MNCs, encompassing various cell types such as lymphocytes (8-10 μm) and monocytes (12-15 μm), exhibit a broader range of speeds (average: 1.89 μm/s in the slowest range (0 - 6 μm/s), increasing to 7.52 μm/s in the middle range (7 - 15 μm/s)) due to this size variation. This heterogeneity in size likely contributes to the wider spread of speeds observed for cells in the control samples containing only MNCs compared to a potentially more uniform cell population like THP-1 cells. In contrast, control samples containing only THP-1 cells, were primarily detecting cells in the faster velocity range (16-40 μm/s) and exhibited a more uniform size distribution with an average diameter ranging from 18 to 21 μm. Blended samples revealed limitations of speed alone for differentiating cells from MNCs in the middle range (7-15 μm/s) due to overlap. However, a key finding emerged as a significant portion of cells within the blended samples occupied a unique higher speed range (16-40 μm/s) absent in control samples containing only MNCs. This suggests these faster cells in the blended samples might be THP-1 cells due to their similar behavior, offering a potential differentiation strategy for further investigation. For instance, the 5% mixture detected >90 cells in the lower speed range, but 11 cells fell within the middle speed range (7-15 μm/s) where both MNCs and THP-1 cells can reside. However, the presence of even a small number of cells (only 2 cells) in the higher speed range (16-40 μm/s) is significant. Since this range is exclusive to THP-1 cells, it highlights the method’s potential to identify 2% (2 out of 100 total mixed cells interrogated) of the 5% THP-1 cells even when the remaining 3% overlap with MNCs in speed at the middle ranges. While the middle speed range warrants further investigation with more runs to definitively characterize the residing cells, the unique higher speed range offers a promising avenue for identifying THP-1 cells, even in the presence of potential confounding factors (instrument limitations, cell health variations, or biological heterogeneity within the cell population) in the middle range. Crucially, the presence of even a single cell in the higher speed range (16-40 μm/s) for the 1% mixture, mimicking a scenario with rare leukemia cells, suggests the method’s potential for identifying as few as 1% of leukemia cells based on their faster movement.

This study presents a label-free, and non-destructive acoustic-based approach for differentiating leukemic cells from healthy cells, offering a non-invasive and potentially faster method compared to traditional bone marrow aspiration. While this study utilized THP-1 leukemia cells, further research is necessary to determine the effectiveness of this approach in differentiating between various normal populations and evolving pre-malignant cells within the same individual. Additionally, challenges such as cost, portability, and throughput need to be addressed before widespread clinical adoption can even be considered. Our research demonstrated distinct movement patterns between THP-1 leukemia cells and normal mononuclear cells (MNCs) within an acoustic field. THP-1 cells exhibited faster movement towards pressure minima (nodes) compared to healthy cells. This difference likely stems from variations in biophysical properties such as density, size, and deformability. Further investigation is necessary to definitively determine the relative contributions of these factors and explore other potential mechanisms, including reasons behind why some THP-1 were observed in the middle range. Studies suggest monocytes, a type of MNC, have a size range of 14-22 μm, a density of approximately 1054 kg/m^3^, and a compressibility of around 400 TPa^−1^. Lymphocytes, another type of MNC, are typically smaller (6-15 μm) and have a similar density (around 1054 kg/m^3^) but slightly lower compressibility (395 TPa^−1^) [8]. THP-1 cells, with a diameter of about 23 μm, are larger than both monocytes and lymphocytes. Based on these observations, we can estimate that THP-1 cells likely have a higher density than 1054 kg/m^3^ and a compressibility exceeding 395 TPa^−1^.

The need for analyzing large cell populations in microfluidic platforms necessitates novel approaches for large-area detection. Our work paves the way for a new generation of such platforms by integrating microfluidics, a synergistic combination of opto-acoustics, and advanced nanofabrication techniques. This approach utilizes piezoelectric ceramic transducers operating at ~1 MHz within a well-characterized detection chamber. Notably, consistent observations across different regions of the chamber suggest the applicability of our method for generating acoustic fields within much larger microfluidic channels, potentially on the scale of millimeters. This represents a significant increase in the analyzed area compared to cell-by-cell flow investigations, allowing us to capture a broader population of cells in a single run (analyzing millions of cells per run). Generating acoustic fields within channels thousands of times larger (millimeters) has the potential to dramatically increase the analyzed area. For instance, transitioning from a 50 μm channel to a 1 mm channel represents a 20-fold increase, while a 5 mm channel offers a staggering 10,000-fold increase. This advancement in large-area detection is particularly beneficial because traditional microscopy techniques often face a trade-off between detailed views and the desired field of view (FOV). For instance, studies investigating rare cells require a large FOV to increase the chance of capturing them, but this might come at the expense of detailed cellular features. We propose addressing this limitation by exploring high numerical aperture objective lens arrays with large area camera sensors, enabling broader, detailed cell visualization with higher magnification power – crucial for capturing rare cells within a larger area. While multi-objective microfluidic chips offer high throughput, they can miss rare cells in unobserved areas. Advances in nanofabrication are key to creating high-resolution, uniform microfluidic structures with improved sensitivity for single cell tracking. This can potentially lead to the analysis of significantly larger cell populations with enhanced detection accuracy.

Furthermore, compact microlenses, phase-contrast illumination, and Light-Emitting Diodes (LEDs) can be integrated into microfluidic chips, enhancing their functionality and expanding their applications [9]. Compact microlenses, fabricated in-situ on microfluidic chips using soft lithography techniques, are designed to focus light beams not only in the horizontal plane but also in the vertical plane. This is achieved by creating a lens chamber with two thin polydimethylsiloxane (PDMS) membranes using the same UV lithography mask as the microfluidic components. Curable polymers are then injected into the lens chamber and cured while pneumatic pressure is applied to keep the PDMS membranes deformed in a quasi-spherical profile. In addition, phase-contrast illumination can be integrated with microfluidic chips. This is done using a polarisation-sensitive camera to simultaneously acquire four images, from which phase contrast images can be calculated. This technique, known as polarisation-resolved differential phase contrast (pDPC) microscopy, can be easily integrated with fluorescence microscopy. Furthermore, OLEDs offer significant advantages for our application due to their ability to achieve sub-millisecond control of illumination [10], [11]. This rapid time resolution allows for precise synchronization of light pulses with acoustic manipulation within the microfluidic channel, a technique known as stroboscopic illumination, potentially enabling us to probe the dynamic behavior of rare cells at high temporal resolution. The well-known advantages of OLEDs, such as homogeneous light distribution over a large surface area and spectrum control, remain crucial for high-throughput analysis and targeted excitation of specific biomarkers in rare cells [10], [12]. These advances not only simplify sample preparation but also eliminate artifacts associated with labels, leading to more accurate analysis of rare cells. To further enhance the platform’s capabilities for rare cell detection in large populations and showcase its flexibility, we propose the integration of microlenses and OLED microarrays directly onto microfluidic chips. Studies discussing the integration of optical detectors, such as silicon photodiode, organic photodiodes (OPDs), and complementary metal–oxide–semiconductor (CMOS) chips, on microfluidic chips support this approach.

Moreover, Surface Acoustic Wave (SAW) devices, known for their ability to generate highly localized fields, can be arranged in multiple arrays. This configuration allows for intricate manipulation strategies, enhancing both sensitivity and throughput. Our study exploits this feature, using large-area microfluidics for cell differentiation. Traditional cell-by-cell flow cytometry methods, which often rely on single-cell interrogation and labeling, may limit throughput and overlook rare cells within a population. However, the multi-array configuration of SAW devices can overcome these limitations. The evolution of microfabrication techniques presents promising solutions for achieving large-area detection and miniaturization, leading to a significant increase in throughput analysis. Microfabrication advancements play a crucial role in achieving high-throughput analysis for large-area detection within our microfluidic platform. One promising approach involves integrating thin-film piezoelectric materials, such as aluminum nitride (AlN) or zinc oxide (ZnO), directly onto microfluidic chips. While these materials offer compelling advantages for miniaturization due to their ability to create “more compact chip designs” [13], [14], their potential impact on large-area detection within the microfluidic channel is also compelling. Advancements in fabrication techniques, such as atomic layer deposition (ALD), can ensure uniform thickness and adhesion of these thin films across a larger surface area [15]. This allows for the generation of acoustic fields across a larger area within the microfluidic chamber, facilitating the manipulation and differentiation of a larger number of cells simultaneously. Additionally, advancements in nanofabrication technologies such as template-assisted assembly or self-assembly methods hold promise for precise alignment and potentially creating micropost arrays or optimized channel geometries. These advancements can further enhance the efficiency of acoustic manipulation within a larger area of the microfluidic channel, ultimately leading to high-throughput analysis of cell populations. Careful engineering efforts are crucial for achieving large-area detection capabilities within our microfluidic platform, with the primary focus on manipulating and analyzing a larger number of cells simultaneously. The state-of-the-art microfabrication techniques such as photolithography and nanoimprint lithography (NIL) could also play a vital role. Photolithography offers a well-established method for patterning large microfluidic channels, while NIL allows for high-throughput replication of these larger features. Additionally, micromachining techniques (e.g. wet etching) can be employed to create specific channel geometries that optimize the generation of acoustic fields across a wider area within the microfluidic channel. Carefully selecting and integrating these techniques is crucial to achieve the optimal design for large-area detection while ensuring compatibility with various cell types, sample volumes, and detection modalities.

## IV. conclusions

This study presents a label-free, and acoustic-based approach for differentiating leukemic cells from healthy cells. Our technology offers a non-invasive and potentially faster method compared to traditional bone marrow aspiration, revolutionizing early-stage leukemia detection with its non-invasive, rapid, and high-throughput capabilities. Leveraging large-area macroscopic detection, as illustrated in Fig. 1, enables the simultaneous observation of individual cell motion for over >10,000 cells per run in under 5 seconds, demonstrating remarkable efficiency and scalability. While flow cytometry remains a mainstay in cell analysis, its reliance on single-cell interrogation and often on labeling can limit throughput and potentially miss rare cells within a population. Our label-free acoustophoretic approach provides a faster and potentially higher-throughput solution for rare cell analysis, offering the potential to analyze larger samples and potentially interrogate individual cells within a population in the next generation of flow cytometry. Beyond leukemia detection, this approach could be harnessed for other cancer risk assessment or even facilitate the label-free detection of airborne particles or microbes.

## V. Acknowledgement

This work was supported by the Biology and Biological Sciences Research Council [grant number BB/X003620/1].

